# Fitness costs in spatially structured environments

**DOI:** 10.1101/012740

**Authors:** F. Débarre

**Affiliations:** Department of Zoology and Biodiversity Research Centre, University of British Columbia, 6270 University Boulevard, Vancouver, B.C., V6T 1Z4, Canada; Centre for Ecology & Conservation, University of Exeter, Penryn Campus, Penryn TR10 9FE, UK

**Keywords:** Fitness costs, Spatial moment equations, Viscous population, Kin Competition, Host defense

## Abstract

The clustering of individuals that results from limited dispersal is a double-edged sword: while it allows for local interactions to be mostly among related individuals, it also results in increased local competition. Here I show that, because they mitigate local competition, fitness costs such as reduced fecundity or reduced survival are less costly in spatially structured environments than in non spatial settings. I first present a simple demographic example to illustrate how spatial structure weakens selection against fitness costs. Then, I illustrate the importance of disentangling the evolution of a trait from the evolution of potential associated costs, using an example taken from a recent study investigating the effect of spatial structure on the evolution of host defense. In this example indeed, the differences between spatial and non-spatial selection gradients are due to differences in the fitness costs, thereby undermining interpretations of the results made in terms of the trait only. This illustrates the need to consider fitness costs as proper traits in both theoretical and empirical studies.

## Introduction

Most populations in nature exhibit some form of spatial structure; this can be because their habitat is fragmented, but also, even in the absence of patchiness, because there are limits to the distances an individual can disperse and because ecological interactions are usually local (Tilman and Kareiva, 1997). Theoretical models have shown that limited dispersal and localised interactions influence demographic processes, such as population growth (Law et al., 2003), epidemiological processes such as the invasion threshold of parasites (Sato et al., 1994; Keeling, 1999), and evolutionary processes, such as the evolution of dispersal (Hamilton and May, 1977; Ferrière and Le Galliard, 2001), altruistic behaviour (Lehmann and Keller, 2006; Lehmann and Rousset, 2010), reproductive effort (Pen, 2000; Lion, 2010), parasite virulence (Boots and Sasaki, 1999; Lion and Boots, 2010) and host defense (Frank, 1998; Best et al., 2011; Débarre et al., 2012; Lion and Gandon, 2015). When dispersal is spatially limited, individuals aggregate within clusters (Lion and van Baalen, 2008) and related individuals tend to live next to one another. This clustering can be beneficial and is for instance key to the evolution of altruism, but it also results in increased local competition, which can annihilate the beneficial effects of clustering (Wilson et al., 1992; Taylor, 1992; Taylor et al., 2011; Débarre et al., 2014). Consequently, traits able to alleviate this local competition can be selected for.

It is usually assumed that new or improved traits come with fitness costs, because of pleiotropic effects or metabolic costs. This is for instance the case for traits of defense against natural enemies. Mounting a defense against parasites can involve the diversion of resources that would otherwise have been used for another purpose (Sheldon and Verhulst, 1996); the chemicals used in the defense can also harm the host (auto-toxicity, Purrington, 2000). In addition to these direct costs, defense traits may also have indirect costs, such as the deterrence of mutualists, or a reduced competitive ability (Strauss et al., 2002). It is therefore common in theoretical studies to assume that the trait of interest is costly. Fitness costs are often considered as a logical necessity to avoid the evolution of Darwinian demons (Reznick et al., 2000) (a situation which, from a theoretical point of view, has a limited interest), but are seldom considered as traits under selection themselves.

Still, when comparing the evolution of a costly trait in spatial *vs.* non-spatial (well-mixed) environments, it is crucial to consider the costs as correlated traits that are also under selection. Indeed, this article shows that spatial structure mitigates fitness costs, and that this result may affect the way we interpret differences in the evolution of specific traits in spatial *vs.* non-spatial contexts, highlighting the limits of verbal interpretations. I first consider a simple model of a population living in a lattice, where reproduction is density-dependent. The decomposition of a selection gradient shows why selection against a reduced fecundity (or against a decreased survival) is less strong in a spatial context than in a non-spatial context. I then assume that individuals can be infected by a directly transmitted parasite, and I study the evolution of reduced susceptibility to the disease, using the same model as Best et al. (2011). Again, I decompose the selection gradient, identifying terms due to the trait itself and terms due to the associated cost, a reduced fecundity. This decomposition reveals that spatial structure does not influence the evolution of reduced susceptibility itself: the change in the evolved level of host susceptibility in spatial *vs.* non-spatial environments is instead a by-product of selection on its associated cost.

## Demographic model

In this first example, we follow the density dynamics of a population of clonally reproducing individuals, when reproduction is density dependent. We assume that there is a large number of breeding sites in the population, and that each site can host at most one individual. Each site is therefore either empty (○) or occupied (*S*).

We denote by *b* individual fecundity (notation is summarized in table 1). An individual can only reproduce if there are empty sites available to host its offspring. Competition between individuals is hence mediated by the presence of accessible empty sites. With probability 1 − *g_R_*, reproduction is local, meaning that the offspring can only be sent to the neighboring breeding sites; with probability *g_R_*, reproduction is global: the offspring can be sent to any empty breeding site in the environment. Death, on the other hand, is density independent, and occurs at a rate *d*; this is the rate at which an occupied site (*S*) becomes empty again (○). We denote by *p_S_* the global density of occupied sites (number of occupied sites divided by the total number of sites in the environment); the global density of empty sites is *p*_○_ = 1 − *p_S_*. The quantity *q*_○|*S*_ is the local density of empty sites around an occupied site. With this, the dynamics of the density of occupied sites can be written using the following spatial moment equation (Rand, 1999; van Baalen, 1998, 2002):

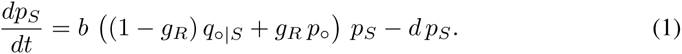

**Table 1:** Notation

|  |  |
| --- | --- |
| $b$ | Fecundity of healthy individuals |
| $d$ | Death rate |
| $\beta$ | Disease transmissibility |
| $\alpha$ | Susceptibility to the disease |
| $\nu$ | Additional death rate due to the disease |
| $g_R$ | Probability of global reproduction |
| $g_T$ | Probability of global transmission |
| $p_x$ | Global density of sites of type $x$ |
| $q_{x\vert y}$ | Local density of sites of type $x$ around a site of type $y$ |
| $\circ$ | Empty site |
| $S$ | Site occupied by an uninfected resident individual |
| $S'$ | Site occupied by an uninfected mutant individual |
| $I$ | Site occupied by an infected resident individual |
| $I'$ | Site occupied by an infected mutant individual |

This population, called the “resident” population, is assumed to be at equilibrium, and we denote by 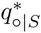 and 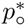 the equilibrium values of the local and global densities of empty sites, respectively. Setting equation (1) equal to zero, we obtain (Lion, 2010)

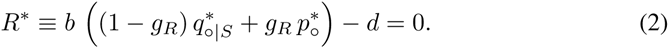

We then assume that a mutant appears, with a different fecundity (*b*′) and/or death rate

(*d*′). The mutant is initially rare, and the invasion dynamics of this rare mutant are given by:

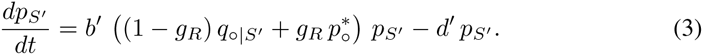

The mutant can establish in the population when *R*′ > 0, with

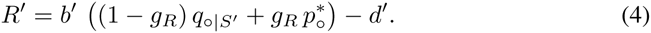

We assume that mutant and resident individuals are phenotypically close: the mutation is of small phenotypic effect, so that we can write *b*′ = *b* + *∂b* and *d*′ = *d* + *∂d*. Consequently, the local density of empty sites seen by a mutant individual is also close to the local density of empty sites seen by a resident individual, so that 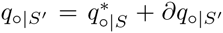. Using the definitions of *R*′ and *R*\* (equations (2) and (4)), we can express the selection gradient *∂R*′ as follows:

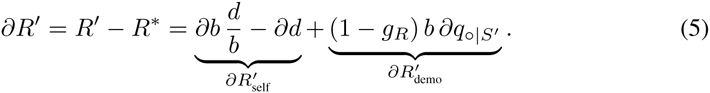

This selection gradient is the sum of two terms. The first term, 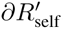, represents the direct effects of the mutation on a mutant’s own fitness; it does not depend on whether reproduction is local or not. The second term, 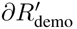, accounts for the changes in the demographic structure of the population due to the mutation, via the term 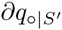, which is the change in the local density of empty sites around an mutant individual, compared to around a resident individual at equilibrium. This second term, proportional to 1 − *g_R_*, vanishes in a non-spatial model, in which *g_R_* = 1.

In this first model, how spatial structure affects the invasion of the mutant is exclusively controlled by 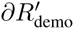, whose sign is the sign of 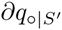. Compared to a non-spatial setting, spatial structure favors the invasion of the mutant if 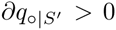, *i.e.*, if mutants see more empty sites around themselves than residents do. This is the crucial point of our argument.

Let us consider a mutant that has a reduced fecundity (*∂b* < 0), a feature that will later be qualified as a fitness cost (the argument goes the same way if we consider changes in the death rate [*∂d* > 0]). In a spatial setting, reproduction is mostly local. Mutants have a lower fecundity, hence have more empty sites in their neighborhood than residents do: 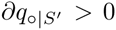, so that 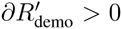. In both cases, though, 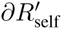 is negative and is the leading term of the selection gradient, so that the mutation is eventually counter-selected. But it is less strongly counter-selected in a spatial context than a a non-spatial context: spatial structure mitigates the fitness cost. Conversely, a mutant with an increased fecundity (*∂b* > 0) sees a lower local density of empty sites 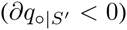, yielding 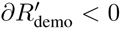; it is therefore less strongly favored in a spatial context than in a non-spatial context. When both fecundity and survival are affected, the sign of 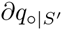 is the sign of *∂d/d* − *∂b/b* (see Supplementary Information): 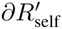 and 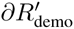 are therefore of opposite signs.

Having more empty sites in its nearest neighborhood 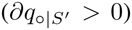 is beneficial to the mutant individual itself, but also to the other individuals that have access to those additional empty sites, potentially other mutants 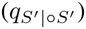: kin competition is reduced.

Figure 1 illustrates the result that fitness costs are less costly in a spatial setting. The selection gradients are calculated numerically, using the pair approximation (Matsuda et al., 1992; Nakamaru et al., 1997) to evaluate local densities. The R codes to run the model as available on figshare, http://dx.doi.org/10.6084/m9.figshare.1183435.

**Figure 1:**
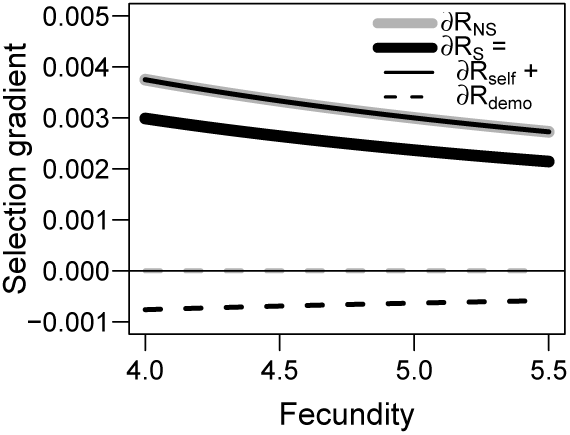
Selection gradients (thick curves) and their decomposition (thin curves) in the demographic model, when only individual fecundity evolves. In grey: selection gradient in a non-spatial model, 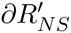 (when *g_R_* = 1); in black, selection gradient in the spatial model, 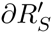 (when *g_R_* = 0). The thin full curve is 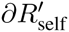; it is the same in both the spatial and non spatial models and appears on top of the thick grey curve; the thin dashed black curve is 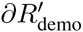. Both *∂R_NS_* and *∂R_S_* are positive: higher values of the fecundity parameter *b* are favored by selection, but *∂R_S_* < *∂R_NS_*. Parameters: *d* = 1, and each individual has *n* = 4 neighbours in the spatial model.

This demographic model has allowed us to show that spatial structure affects the magnitude of the effect of the fitness cost: it makes fitness costs less costly. We will now see why this matters.

## Evolution of host susceptibility

We now consider the evolution of a trait of defense against parasites, namely, the evolution of avoidance (or reduced susceptibility), as studied by Best et al. (2011). I use the same model and the same assumptions as Best et al., but with the notation of Débarre et al. (2012), where the decomposition of the selection gradient used in this study was introduced. As in Best et al. (2011), we will assume that defense is costly, and that a reduced susceptibility to the infection comes at the cost of a reduced fecundity.

The basic assumptions are the same as previously (one individual per site, density-dependent reproduction), but we now assume that the individuals can be infected by a parasite. Infected individuals cannot reproduce nor recover: the infected state is a dead-end, and we denote by *ν* the additional mortality due to the infection (also called virulence (Read, 1994)). With a probability 1 − *g_T_*, an infected individual can only infect its (healthy) neighbors; with probability *g_T_*, transmission is global: an infected individual can infect any healthy individual in the population. A parameter *β* denotes the transmissibility of the parasite, while a parameter *α* denotes the susceptibility of a healthy host. With these assumptions, the dynamics of the density of sites occupied by healthy (*p_S_*) and infected (*p_I_*) individuals are given by the following system (notation is recapitulated in table 1):

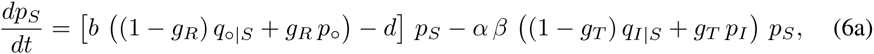

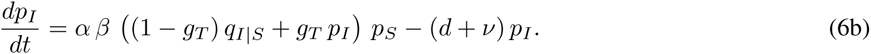

As previously, we assume that the population (called the “resident” population) is at equilibrium and we use a star * to denote global and local densities evaluated at this equilibrium. We assume that a mutant appears, with a different susceptibility to the infection *α*′ = *α* + *∂α*, and different fecundity, *b*′ = *b* + *∂b* (the product *∂α.∂b* is positive). The sign of the selection gradient *∂R*′ indicates whether these mutants can establish; Débarre et al. (2012) have shown that the selection gradient can be expressed as follows:

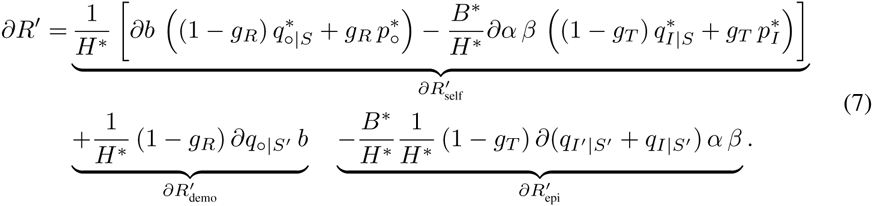

where

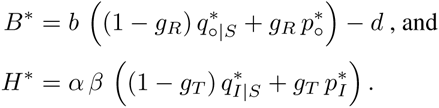

The method to derive equation (7) is detailed in Débarre et al. (2012, Appendix C); equation (7) can be further simplified by noting that *B*\* = *H*\*. We also note that the expression of *B*\* is identical to the expression of *R*\* in the demographic model (equation (2)), except that this quantity is not equal to zero anymore, for the density of healthy individuals is also affected by infection dynamics (see equation (6a)).

The interpretation of the first two terms of the selection gradient (7) is the same as in the previous section: 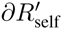, corresponds to the direct effects of the mutation on the mutants’ own fitness, and 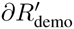 takes into account changes in the demographic structure of the population. A third term, 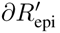, corresponds to changes in the epidemiological structure of the population, via the terms 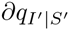 and 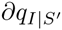, whose sum corresponds to the changes in the density of infected individuals (resident or mutant) in the neighborhood of a healthy mutant individual. Both 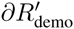 and 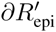 vanish in a non-spatial context, when reproduction and transmission are purely global (*g_R_* = *g_T_* = 1).

The selection gradient (7) conflates the effects of changes in the trait of interest (*α*) and the associated cost (*b*), but we can disentangle these effects, by noting that

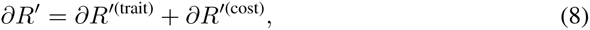

where *∂R*′^(trait)^ is the selection gradient that we would obtain if the trait under selection had no associated cost (*b*′ = *b*), while *∂R*′^(cost)^ is the selection gradient obtained when mutants only carry the cost, but have the same trait as the residents (*α*′ = *α*). Both can be further subdivided into direct, demographic and epidemiological components, as in equation (7).

Let us compare the global selection gradient in a purely spatial setting (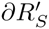, when *g_R_* = *g_T_* = 0) and purely non-spatial setting (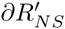, when *g_R_* = *g_T_* = 1)—intermediate values of *g_R_* and *g_T_* yield results that are intermediate between these two extremes. Figure 2(a) shows that 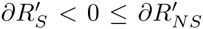: lower susceptibility to the disease (an avoidance defense mechanism) evolves in a spatially structured environment, while a non-spatial environment selects for a higher susceptibility to the disease. A dissection of the selection gradient is going to tell us where this difference comes from.

**Figure 2:**
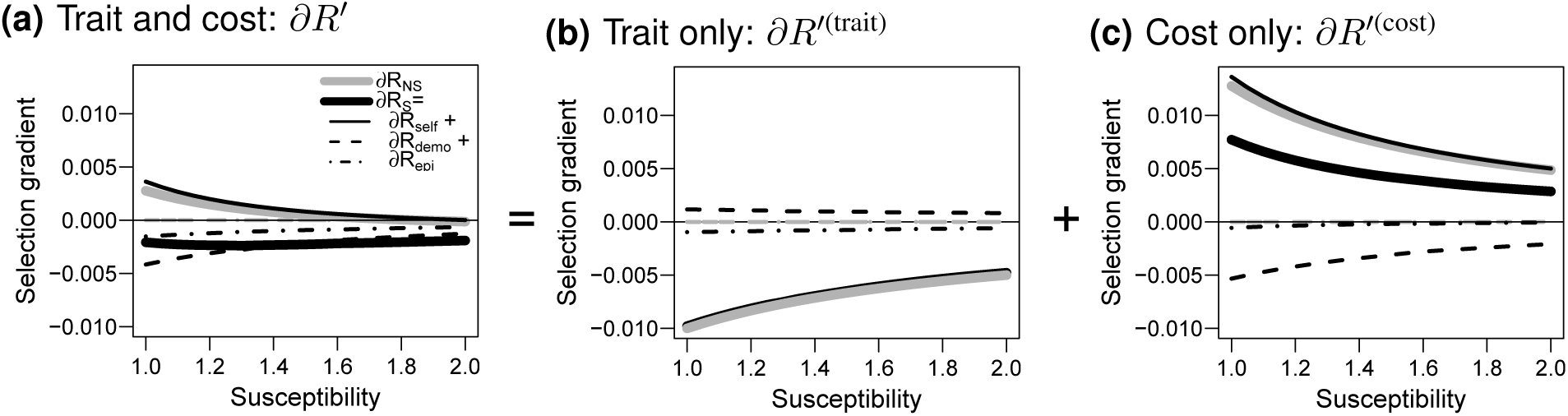
Selection gradients (thick curves) and their decomposition (thin curves), in the non-spatial (grey; *g_R_* = *g_T_* = 1) and spatial (black; *g_R_* = *g_T_* = 0) models. In (a), a mutation affects both the trait (susceptibility to the disease, *α*) and the cost (fecundity, *b*). In (b), only the trait evolves, mutants have the same fecundity as residents; in (c), only the cost evolves, mutants have the same susceptibility as residents. Parameters: same as in figure 2a in Best et al. (2011): *d* = 0.1, *β* = 1, *ν* = 0.1, *b*(*α*) = 4*(−0.2+1.2*α*)/(0.9+0.1*α*), and *n* = 4 neighbours in the spatial model.

Let us start with the effect of the cost, a reduced fecundity, because this effect is similar to the situation studied in the demographic model (see figure 2(c)), except that the 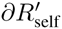 terms now differ between the spatial and non-spatial settings (in figure 2(c), the thick grey curve and thin black curve are not exactly superimposed anymore), and there is an additional 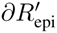 term in the spatial setting (dot-dashed curve). The argument however remains: the cost is less costly in a spatial setting.

Let us now turn to the effect of the trait itself, in the absence of cost, and consider mutant individuals that are less susceptible to the disease. In a spatial context, mutants are also more likely to be surrounded by other mutants than by residents; hence, mutants are less likely to have infected neighbors than residents do. Kin selection accounts for why the epidemiological component of spatial structure 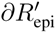 favors lower susceptibility (see the dot-dashed curve in figure 2(b)). However, a lower susceptibility has also demographic consequences. Being less susceptible to the disease means being less likely to die because of it, and being less likely to be castrated by the parasite. Hence, mutants have comparatively fewer empty sites in their neighborhood: kin competition is higher. This is why the demographic component 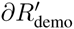 favors higher susceptibilities to the disease instead (see the dashed curve in figure 2(b)). Interestingly, in the spatial context, the effects of the demographic and epidemiological structures almost compensate each other. As a result, 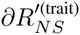 and 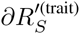 are almost identical: the thick grey and thick black curves are almost on top of each other in figure 2(b). The overall effect of spatial structure on the evolution of susceptibility to the disease is therefore almost negligible, but it is interesting to note that it is such that spatial structure leads to a slightly weaker selection for lower susceptibility—the opposite of the conclusion drawn when a fecundity cost is associated.

Now going back to the global selection gradient, encompassing the trait and its associated cost (figure 2(a)), we now understand that the difference between the spatial and non-spatial settings are in fact mostly driven by the fitness cost, and not by the trait of interest itself.

This conclusion is drawn using the same parameters as Best et al. (2011), but does it hold for other combinations of parameters? To answer this question, I drew random sets of parameters and computed the selection gradients. The resulting distributions of selection gradients in non-spatial and spatial settings, as well as the distribution of their difference, are represented in figure 3. Overall, the tested random combinations of parameters confirm our conclusions: when the defense trait is costly, selection is more towards a lower susceptibility in a spatial setting than in a non-spatial one (figure 3(a)); the cost is less costly in a spatial setting (figure 3(c)); finally, selection on the susceptibility trait is of similar magnitude in spatial and non-spatial settings, and is slightly towards higher susceptibility in a spatial setting (figure 3(b)).

**Figure 3:**
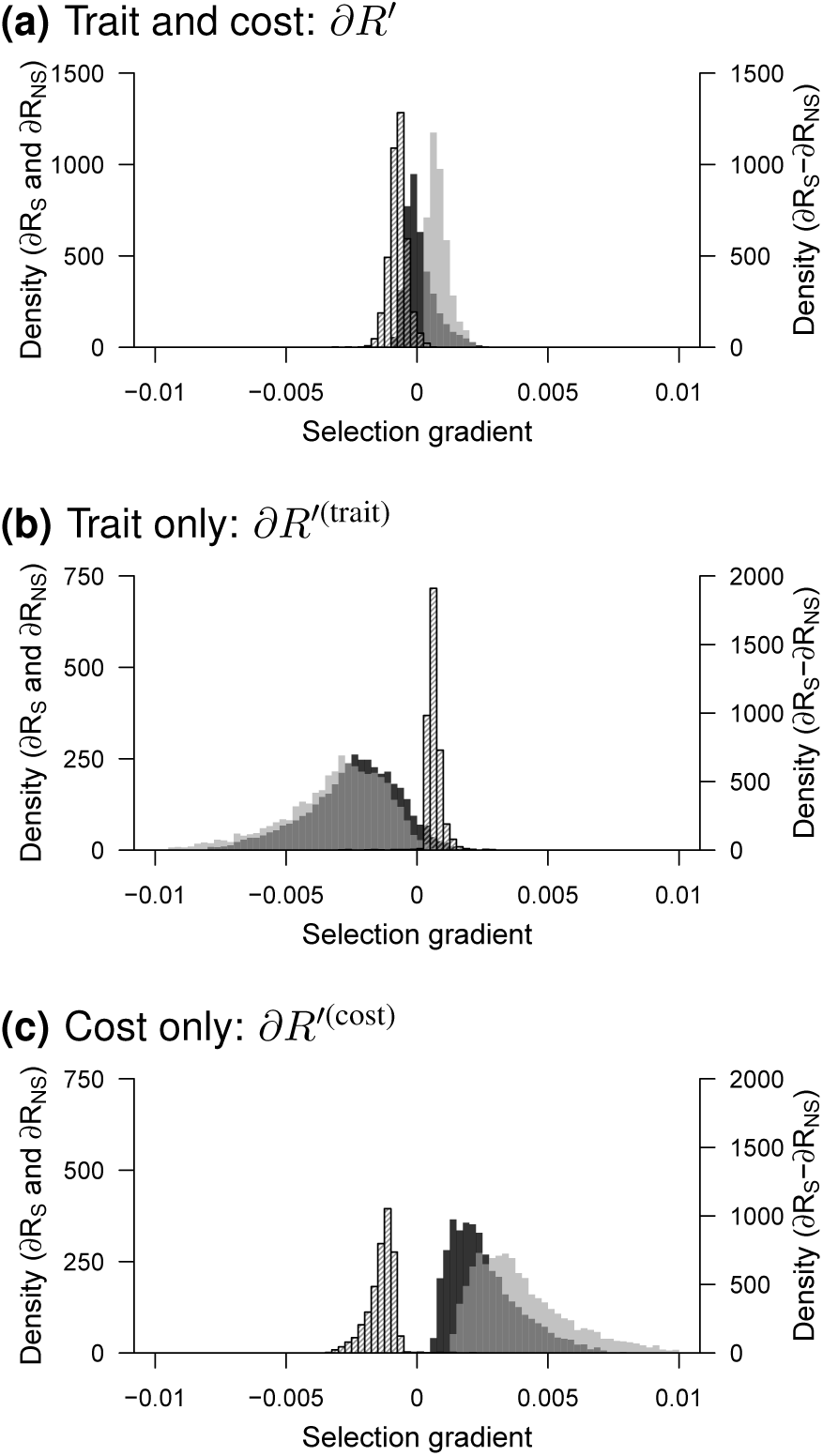
Distributions of values of the selection gradient in fully spatial (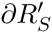, black) and non-spatial (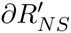, gray) models, and of their difference (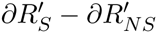, hatched) for different combinations of traits. In (a), a mutation affects both the trait (susceptibility to the disease, *α*) and the cost (fecundity *b*). In (b), only the trait is affected, and in (c) just the cost is affected. The different traits (6000 random combinations) were independently drawn from uniform distributions: *α* ∈ {1, 3}, *b* ∈ {3, 9}, *d* ∈ {0.05, 0.15}, *ν* ∈ {0.01, 0.15}; *β* = 1.

## Discussion

While evolutionary studies commonly assume that a change in trait of interest comes with an associated cost, the cost itself is seldom considered as a trait in its own right. In this article, I show the importance of disentangling the effects of a change in a trait of interest from the effects of a change in the associated cost. To illustrate this point, I use an example taken from a recent study by Best et al. (2011), investigating the effect of spatial structure on the evolution of host susceptibility to a disease. Assuming that a lower susceptibility to a disease is associated to a lower host fecundity, Best et al. (2011) found that higher levels of host susceptibility evolve in a non-spatial setting than in a spatial setting. Decomposing selection gradients into terms due to the trait (host susceptibility) and the cost (fecundity), and dissecting these terms into a direct effect (effect of the change on the individual itself), as well as demographic and epidemiological effects (changes in the spatial structure of the population), I show that spatial structure actually almost does not influence the evolution of host susceptibility strictly speaking. Instead, spatial structure makes fitness costs less costly, which indirectly leads to lower levels of host susceptibility to the disease in a spatial setting.

Why are fitness costs less costly in a spatial setting? Let us go back to the demographic model (without infected individuals) to explain this result. The model assumes that reproduction is density-dependent: there is a fixed (but large) number of breeding sites, and an individual can only reproduce if it has access to empty sites. When reproduction is local, an individual can only reproduce in the nearby sites; this is why the quantity 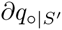, representing the difference in the density of neighboring empty sites near a mutant *vs.* a resident, is key. A mutant which reproduces less will have more empty sites nearby 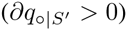: local competition is reduced for mutants. In a non-spatial setting (*g_R_* = 1), where an individual can send its offspring to any empty site in the entire environment, mutants and residents have access to the same number of empty sites, so there is no beneficial effect of a lower reproduction or survival. In both cases, a lower fecundity remains directly detrimental, and is therefore not selected for, but selection against a lower fecundity is weaker in a spatial setting (see figure 1).

And yet, in a seminal model, Frank (1998) showed that increased relatedness led to more disease avoidance, equivalent to a lower host susceptibility. How does Frank’s result compare to ours? Importantly, Frank’s kin selection model lacks two features that are key to our model. First, there are not any explicit epidemiological dynamics in Frank’s model: the probability of future attack (parameter *a* in his model) is a constant. In other words, the force of infection is constant, meaning that the dynamics of parasite densities are not affected by the availability of hosts in the population: there are no epidemiological feedbacks (Boots et al., 2009). Second, and more importantly, there is no density-dependence in Frank’s model: the fitness of individuals does not depend on the local or global density of hosts—in other words, there are no demographic feedbacks. The absence of density-dependence has two consequences. First, the fitness cost (*c* in Frank’s model) is not less costly in a spatial setting, because there is no local competition for space that the cost could alleviate. Second, the demographic term of the selection gradient, that compensated the epidemiological term, disappears, as if we removed the dashed curve in figure 2(b): the selection gradient in the spatial setting changes and becomes in favor of a lower susceptibility.

To conclude, our study shows that, when reproduction is density-dependent, fitness costs are less costly in a spatial setting than in a non-spatial setting. This highlights the need to consider costs as correlated traits in both empirical and theoretical studies, and whenever possible, to study the effects of the trait and the costs independently, to disentangle their relative contributions. On a positive note finally, the finding that spatial structure actually almost does not directly influence the evolution of host susceptibility undermines Best et al. (2011)’s dramatic conclusion that a more globalized world could lead to lower levels of host defense against parasites.

## Acknowledgements

I thank Sébastien Lion, François Blanquart and Mike Boots for comments on previous versions of this work, as well as Jon Wilkins and two anonymous reviewers for useful suggestions. Funding was provided by an NSF grant DMS 0540392, a Natural Sciences and Engineering Research Council CREATE Training Program in Biodiversity, and a fellowship of the Wissenschaftskolleg zu Berlin.

## Supplementary Information

### A Demographic model: finding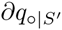

### A.1 Derivation using the full model

A first expression of the selection gradient is given in the main text, equation (5); it was obtained from the dynamics of the global densities. The aim of this section is to derive another expression of the selection gradient, this time using pair dynamics, to eventually derive an expression for 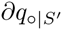. This section describes the steps to obtain the result; more details can be found in a supplementary Mathematica file available on Figshare, http://dx.doi.org/10.6084/m9.figshare.1183435.

#### A.1.1 Pair dynamics

We denote by **p** the column vector 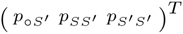 of pairs of neighboring sites inhabited by mutant individuals, and by **q** the corresponding vector of local densities 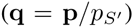. The symbol *n* representing the number of neighbors, we define *ϕ* = 1/*n* and 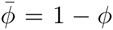. The dynamics of **p** can be written as

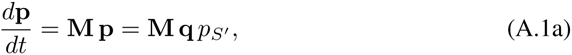

where

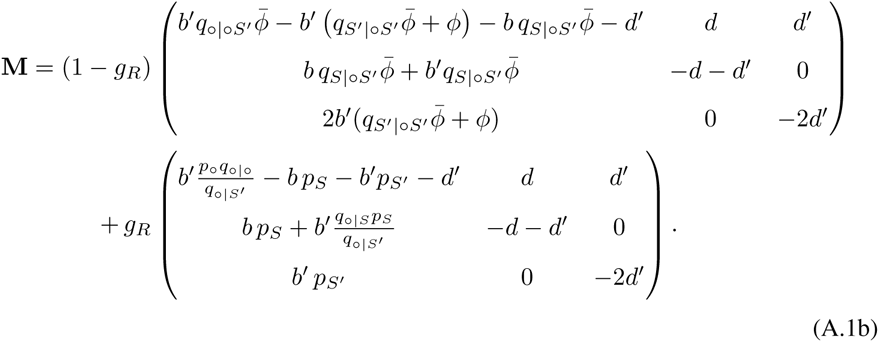

We are interested in the initial dynamics of the mutants, so we evaluate the matrix **M** when 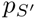 is close to zero and *p_S_* is at the equilibrium in the absence of mutants 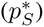.Then, we use the fact that the mutation is of small effect, so that we can write the local densities as follows:

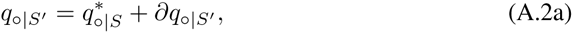

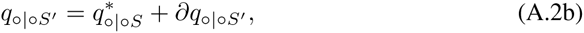

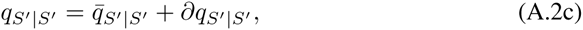

where 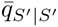 is the nearest neighbor relatedness with rare mutants, the bar 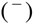 meaning that the mutation is neutral (Lion and Gandon, 2009; Lion, 2010). We also note that

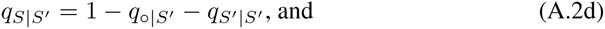

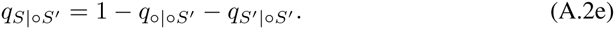

We can therefore approximate the matrix **M** as a sum of two matrices **M_0_** and **M_1_**, where **M_0_** contains the terms corresponding to a neutral mutant, while **M_1_** contains all the *∂* terms.

While the dynamics of the global frequencies are slow (involving *∂* terms only), the dynamics of the pairs occur at another time scale (**M** is *O*(1)). The local densities therefore equilibrate before the global frequencies, as is the case in spatial models when selection is weak (Matsuda et al., 1992; van Baalen and Rand, 1998; Lion and van Baalen, 2009). The vector of local densities **q** reaches a quasi-equilibrium 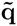 and the matrix **M** converges towards a constant matrix 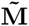. The dynamics of 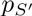 can then be rewritten as 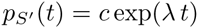, where *λ* is the dominant eigenvalue of 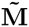 (van Baalen, 1998; Ferrière and Le Galliard, 2001).

#### A.1.2 Evaluation of the local densities for a neutral mutant

The first step is to obtain 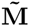 is to find the local densities of the resident (*) and neutral mutant 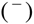. At neutrality (the mutant’s trait values being equal to the resident’s trait values), once the quasi-equilibrium is reached, we have

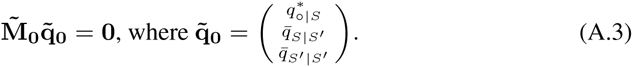

From equation (2) in the main text, we have (when *g_R_* > 0)

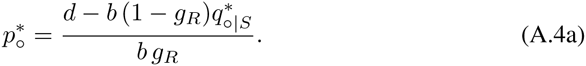

Solving (A.3), we find the following equalities:

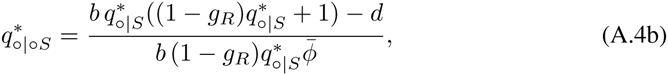

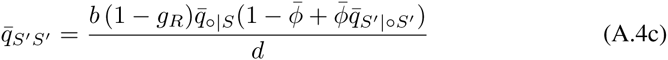

We also note that the vector **v_0_** = (2 1 1) is a left eigenvector of 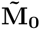, associated to the eigenvalue 0. We therefore have 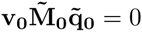.

#### A.1.3 Leading eigenvalue of 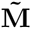

We now want to find the leading eigenvalue of 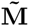. With a non-neutral mutant, 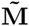 differs from 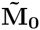 by a perturbation 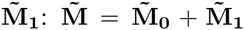. We can therefore approximate the leading eigenvalue of 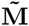 as (Lion, 2010)

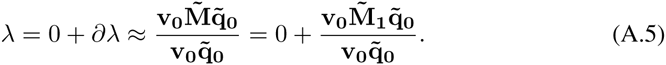

We have already found the invasion exponent of the system, which we call *∂R*′ (equation (5)); by solving the equation *∂λ* = *∂R*′ for 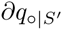, we obtain

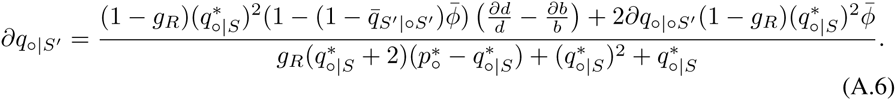

Neglecting the effect of the mutation on the presence of empty sites connected to neighboring empty sites (that is, setting 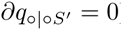), equation (A.6) reduces to

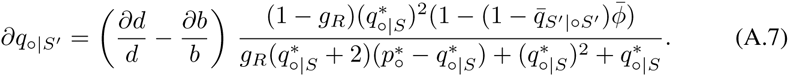

In the next part, we will confirm that 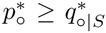, so that the sign of 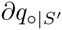 is given by the sign of 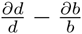.

We recall that our numerical simulations use the pair approximation (PA) (Matsuda et al., 1992; Nakamaru et al., 1997): that is, we approximated quantities like *q_a_*_|*bc*_ as *q_a_*_|*b*_. This technique allows us to have as many equations as there are variables in the system (*i.e.*, to close the system), and to numerically solve it. Technically, this means that in the simulations, 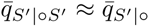, which is close to zero when the mutant is rare. In other words, in the simulations,

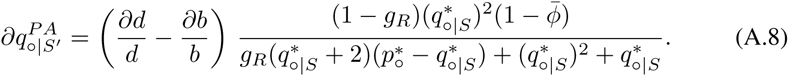

### A.2 Resident dynamics with the pair approximation

In this section, we use the pair approximation to show that 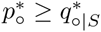.

We consider the system of pair dynamics in the resident population (*i.e.*, the equivalent of (A.1) with the residents only). With the pair approximation, we can simplify both 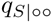 and 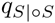 into 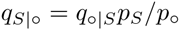. At the equilibrium, we find that 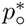 and 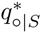 must satisfy

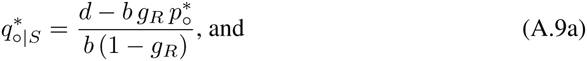

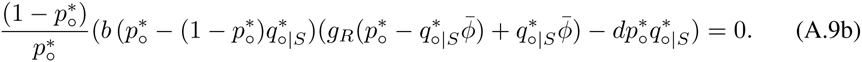

We want to show that 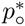, which is root of (A.9b), is greater or equal than *d/b*.

In the fully spatial model (*g_R_* = 0), we find that (recall that 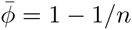)

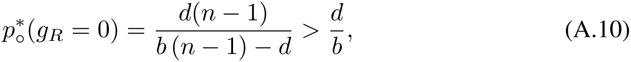

while in the non-spatial model (*g_R_* = 1), we have

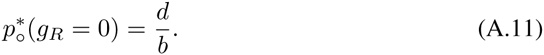

By continuity, if 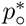 goes below *d/b* for intermediate values of *gR*, then it must go through *d/b*. So we can search when *d/b* is a root of (A.9b). We find that this is the case only when *g_R_* = 1 or 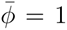 (which means that there is an infinite number of neighbors, and is therefore the same as having *g_R_* = 1). Because 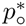 does not go through *d/b* when 0 < *g_R_* < 1, then it remains above it.

From (A.9a), 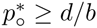 implies that 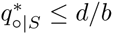, so that we have 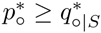.

